# Inhibition of Mitochondrial Fission and iNOS in the Dorsal Vagal Complex Protects from Overeating and Weight Gain

**DOI:** 10.1101/2020.06.26.173641

**Authors:** Bianca Patel, Lauryn New, Joanne C. Griffiths, Jim Deuchars, Beatrice M. Filippi

## Abstract

The dorsal vagal complex (DVC) senses changes in insulin levels and controls glucose homeostasis, feeding behaviour and body weight. Three days of high-fat diet (HFD) in rats is sufficient to induce insulin resistance in the DVC and impair its ability to regulate feeding behaviour. HFD-feeding is associated with increased mitochondrial fission in the DVC and fission is regulated by dynamin-related protein 1 (Drp1). Higher Drp1 activity can inhibit insulin signalling, although the exact mechanisms controlling body weight remain elusive. Here we show that Drp1 activation in DVC leads to higher body weight in rats and Drp1 inhibition in HFD-fed rats reduced body weight gain, cumulative food intake and adipose tissue, and prevented insulin resistance. Rats expressing active Drp1 in the DVC had higher levels of inducible nitric oxide synthase (iNOS) and knockdown of iNOS in the DVC of HFD-fed rats led to a reduction in body weight gain, cumulative food intake and adipose tissue, and prevented insulin resistance. In obese insulin-resistant animals, inhibition of mitochondrial fission or DVC iNOS knockdown restored insulin sensitivity and decreased food intake, body weight and fat deposition. Finally, we show that inhibiting mitochondrial fission in DVC astrocytes is sufficient to protect rats from developing HFD-dependent insulin resistance, hyperphagia, body weight gain and fat deposition. Our study uncovers new molecular and cellular targets for brain regulation of whole-body metabolism, which could inform new strategies to combat obesity and diabetes.

## Introduction

Diabetes and obesity are epidemic diseases with rising incidence across the world. Overnutrition is the predominant pathogenic inducer of insulin resistance, which is mainly caused by increased circulating levels of glucose, free fatty acids and amino acids (Dresner *et al*, 1999; Roden *et al*, 1996). Increase of these circulating factors causes mitochondrial oxidative stress and endoplasmic reticulum (ER) stress in peripheral tissues (Birkenfeld *et al*, 2011) and the central nervous system (CNS) (De Souza *et al*, 2005). The CNS is instrumental in regulating metabolic homeostasis as it receives and processes peripheral inputs reporting the metabolic status of an individual and signals back to peripheral organs to maintain energy balance. Subtle imbalance of these homeostatic processes can lead to metabolic diseases with obesity and diabetes causing a major human health problem and a strain in health services worldwide. Overnutrition has a large impact on the CNS and leads to a loss of the brain’s ability to sense changes in hormone and nutrient levels. This has a dramatic effect on metabolic homeostasis where individuals struggle to maintain whole-body energy balance, leading to obesity and the development of diabetes (Timper & Brüning, 2017; Cai, 2013). Restoring the brain’s ability to modulate metabolic functions could be very important to prevent the negative outcomes of obesity and diabetes.

Insulin is one of the major humoral signals which modulates energy balance. Deletion of the insulin receptor in the brain (NIR knockout mice) led to increased body fat and insulin resistance (Bruning *et al*, 2000). The hypothalamus, which is located at the lateral boundaries of the third ventricle, is one of the major areas of the brain responsible for regulating energy homeostasis in response to changes in hormonal and nutrient levels (Timper & Brüning, 2017). Intracerebroventricular (ICV) injection of insulin in the mediobasal hypothalamus (MBH) decreased hepatic glucose production (HGP) (Obici *et al*, 2002b; Pocai *et al*, 2005), while selective decrease in insulin receptor levels in the hypothalamus resulted in hyperphagia and increased fat mass (Obici *et al*, 2002a). In rodents, the dorsal vagal complex (DVC) of the brain is another important regulator of glucose metabolism (Filippi *et al*, 2012) and food intake (Filippi *et al*, 2014). In particular, the nucleus of the solitary tract (NTS) in the DVC senses insulin and triggers a neuronal relay to decrease HGP and food intake (Filippi *et al*, 2012; Filippi *et al*, 2014). Interestingly, directly targeting the brain by intranasal delivery of insulin lowers blood glucose levels and decreases food intake in non-obese and non-diabetic humans (Dash *et al*, 2015; Hallschmid *et al*, 2004). This suggests that insulin action in the brain may be clinically relevant for treatment of obese and diabetic individuals, and mandates further studies to fully understand the mechanisms of brain insulin signalling in animal models of obesity and diabetes.

Rodents lose the ability to sense both MBH and DVC insulin, and regulate HGP and food intake (Filippi *et al*, 2012; 2014;Ono *et al*, 2008) after just 3-days of high-fat diet (HFD) feeding. The development of insulin resistance in the brain was associated with increased inflammation and ER stress (Cai & Khor, 2019; Zhang *et al*, 2008). HFD, obesity and insulin resistance in the brain have also been associated with mitochondrial dysfunction (Nasrallah & Horvath, 2014; Jin & Diano, 2018). Mitochondria are highly dynamic organelles that change morphology via fission and fusion to meet energy demands (Jheng *et al*, 2012). When cellular energy demands are high, mitochondrial fusion is one way to produce more ATP (Gao *et al*, 2014; Chan, 2006). Conversely, when the cell is in excess of energy, there is an increase in mitochondrial fragmentation (fission) that causes decreased mitochondrial activity and mitophagy (Gao *et al*, 2014; Chan, 2006). These two opposing processes are regulated by several proteins, mitochondrial fusion is regulated by mitofusin 1 and 2 (Mfn-1 and −2) and optic atrophy (Opa1), while mitochondrial fission is regulated by dynamin-related protein 1 (Drp1) and fission protein 1 (Fis1); a fine balance of these processes is needed to maintain mitochondria functionality (Gao *et al*, 2014). In adipose tissue, mitochondrial dysfunction increases oxidative stress, leading to an increase in fat oxidation and lipid accumulation which is associated with insulin resistance (Gao *et al*, 2014). In particular, enhanced mitochondrial fission has been associated with diet-induced obesity and insulin resistance in different tissues. An increase in active Drp1 caused high levels of mitochondrial fission in skeletal muscle and this was associated with diet-induced obesity and insulin resistance (Jheng *et al*, 2012). In addition, deletion of Drp1 in the liver prevents mice from developing diet-induced insulin resistance (Wang *et al*, 2015). Mitochondrial dynamics also play a pivotal role in how neuronal cells respond to hormonal changes. In POMC and AgRP neurones the knockdown of proteins involved in the regulation of mitochondrial dynamics can alter feeding behaviour and glucose metabolism (Schneeberger *et al*, 2013; Jin & Diano, 2018; Santoro *et al*, 2017; Dietrich *et al*, 2013), while the glucose-sensing neurones of the ventromedial hypothalamus change mitochondrial morphology and their firing rates in response to changes in glucose levels (Toda *et al*, 2016). Diet-induced alteration of mitochondrial dynamics also occurs in extra hypothalamic regions. Indeed, HFD-dependent insulin resistance in the DVC is caused by increased mitochondrial fission through activation of Drp1 (Filippi *et al*, 2017). Direct injection of chemical or molecular inhibitors of Drp1 into the DVC restored the ability of insulin to regulate glucose metabolism in HFD-fed rats (Filippi *et al*, 2017). Conversely, direct injection of an adenovirus expressing active Drp1 in the DVC caused insulin resistance in rats fed with regular chow (RC) diet (Filippi *et al*, 2017) and prevented DVC insulin from lowering HGP. However, little is known whether the changes in mitochondrial fission in DVC neurones affects food intake and body weight.

This is an important biological question as it is not yet clear how HFD-dependent activation of Drp1 and mitochondrial fission cause insulin resistance. Interestingly however, there is an increase in inducible nitric oxide synthase (iNOS) levels in the DVC of HFD-fed rats and of RC-fed rats overexpressing active Drp1 (Filippi *et al*, 2017). Studies in rodents suggest that increased iNOS levels cause insulin resistance in muscle of diet-induced obese and genetically obese mice (Carvalho-Filho *et al*, 2005), while S-nitrosylation and activation of ER stress-related proteins in the liver of HFD-fed rats caused alteration of metabolic functions (Yang *et al*, 2015). Furthermore, in the hypothalamus, increased iNOS levels triggered insulin resistance and obesity (Katashima *et al*, 2017). Therefore, we investigated whether changes in iNOS levels in the DVC can affect food intake and body weight.

Aberrant levels of iNOS in astrocytes leads to astrogliosis and has been shown to increase levels of neuroinflammation and neurotoxicity (Liberatore *et al*, 1999), suggesting that astrocytes maybe involved in the link between HFD induced elevation in iNOS levels and insulin resistance. Hypothalamic astrocyte-dependent inflammation (astrogliosis) was seen in both diet-induced or genetically modified models of obesity, however the exact mechanism of this is not well understood (Buckman *et al*, 2015; Horvath *et al*, 2010). Whether changes in mitochondrial dynamics and iNOS levels in astrocytes is involved in the development of DVC-insulin resistance, needs further investigation.

Here we show that changes in mitochondrial dynamics in the NTS of the DVC affect insulin sensitivity, food intake, body weight and fat deposition. We demonstrate that increasing mitochondrial fission by activating Drp1 in the NTS causes insulin resistance, hyperphagia and body weight gain. Conversely, inhibiting Drp1 to decrease mitochondrial fission, protects from HFD-depended insulin resistance and decreases food intake and body weight gain. In addition, we show that HFD and activation of Drp1 causes an increase in iNOS levels in the DVC and it is sufficient to knock down iNOS to protect from HFD-dependent development of insulin resistance. Finally, for the first time we show that it is sufficient to inhibit mitochondrial fission in astrocytes of the NTS to protect rodents from developing HFD-dependent insulin resistance, as well resulting in a decreased food intake, body weight gain and fat deposition.

The brain is the central player in maintaining energy balance, alteration of the brain’s ability to keep metabolic homeostasis is strictly correlated with the development of obesity. A better understanding of the brain regions involved controlling metabolic functions, and of the molecular mechanisms that alters these functions, is at the basis for developing new strategies to fight the obesity epidemic

## Results

### Higher mitochondrial fission in the DVC causes insulin resistance, hyperphagia, and body weight gain

The DVC senses insulin and decreases food intake, this effect is lost in HFD-fed, insulin resistant rats. Increased mitochondrial fission was observed in the DVC of HFD-fed rats (Filippi *et al*, 2017) and we sought to determine whether HFD-dependent increased mitochondrial fission in the DVC affects the ability of insulin to lower food intake. In addition, we aimed to determine whether chronic alteration of mitochondrial dynamics in the DVC can affect the feeding behaviour of rats. Using an adenoviral system, we expressed a CMV-driven constitutively active form of Drp1 (Drp1-S673A) in the NTS of the DVC which causes an increase in mitochondrial fission (Filippi *et al*, 2017). A GFP expressing adenovirus was used as control (Fig 1 A and B). Following viral injection, we monitored food intake and body weight for 2 weeks (Fig. 1A). To determine whether changes in mitochondrial fission can affect the ability of DVC-insulin to decrease food intake, on week 1 and 2 we performed an acute feeding study where insulin was injected in the DVC (targeting the NTS), and food intake was monitored for 4 hours (Fig. 1A). Rats expressing Drp1-S637A were insulin resistant, hyperphagic, had increased body weight and accumulated more fat compared to the GFP-expressing control rats (Fig.1C to F). A single injection of insulin in the DVC was sufficient to decrease food intake in regular chow-fed rats within 4 hours. Expression of the active form of Drp1 in the DVC induced insulin resistance and impaired insulin-dependent decrease in food intake in RC-fed rats (Fig. 1C). In addition, chronic activation of Drp1 in the DVC was sufficient to induce hyperphagia (Fig. 1D), body weight increase (Fig. 1E) and increase in total white adipose tissue accumulation (sum of retroperitoneal, epididymal and visceral fat) (Fig. 1F). These data show that increasing Drp1-dependent mitochondrial fission by constitutive activation of Drp1 in the DVC is sufficient to cause insulin resistance and increase food intake and body weight gain.

**Figure 1:**
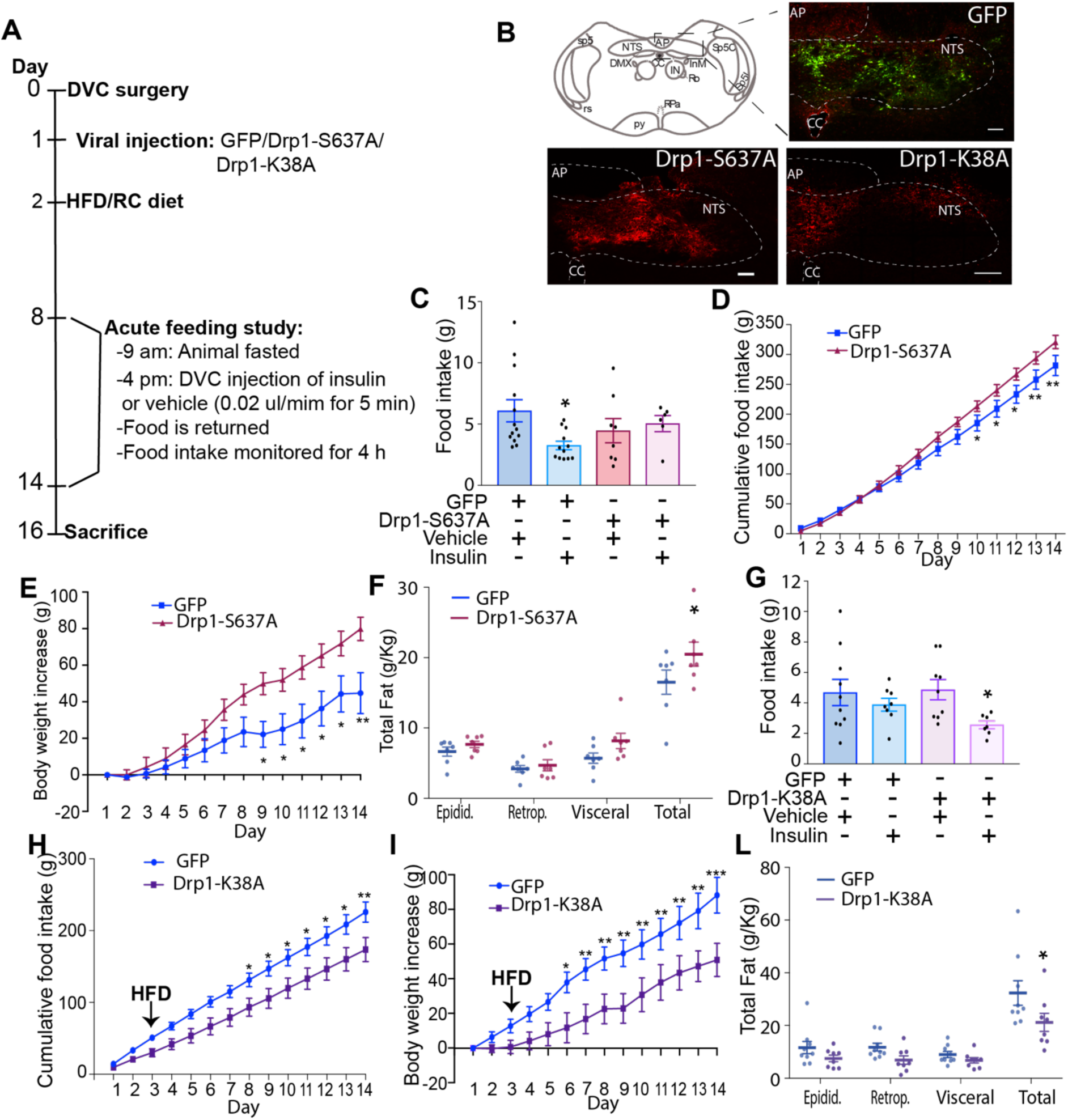
Changes in mitochondrial fission in the DVC affect insulin sensitivity, food intake, body weight and fat deposition. (**A**) The experimental design including feeding study procedure. (**B**) Representative confocal image of DVC areas expressing FLAG-tagged mutants of Drp1 (constitutively active Drp1-S637A and dominant negative Drp1-K38A) or GFP, in the NTS of the DVC. Scale bar=100 μm. (**C and G**) Acute feeding study: animals fed with RC (C) or HFD (G) were fasted for 7 hours and then infused bilaterally into the DVC with a total 0.2 μl of insulin or a vehicle over 5 minutes. Food was then returned and food intake was observed every half hour for 4 hours. Data are shown as mean ± SEM, with each single point highlighted. In C, data are representative of n=13 for GFP vehicle, n=12 for GFP insulin, n=8 for Drp1-S637A vehicle, n=6 for Drp1-S637A insulin. In G, data are representative of n=10 for GFP vehicle, n=8 for GFP insulin, n=9 for Drp1-S637A vehicle, n=7 for Drp1-S637A insulin. (**D and H**) Cumulative food intake from day 1. (**E and I**) Body weight increase from day 1. (**F and L**) White adipose tissue measurements: epididymal, retroperitoneal and visceral fat collected on the day of sacrifice. Data are shown mean ± SEM, with each single point highlighted. Data are representative of n=7 rats for both GFP and Drp1-S637A in D to F and n=9 rats for GFP and n=8 rats for Drp1-K38A in H to L.*p < 0.05, **p <0.01, *** p<0.001.

### Inhibition of mitochondrial fission in the DVC protects from developing HFD-dependent insulin resistance and decreases body weight and food intake

Three days of HFD-feeding are sufficient to cause insulin resistance in the DVC and prevent an insulin-dependent decrease in food intake. There is also a clear increase in mitochondrial fission following 3 days of HFD-feeding (Filippi *et al*, 2017), therefore we wanted to determine whether reducing mitochondrial fission could protect rats from losing insulin sensitivity and affect feeding behaviour. Inhibition of mitochondrial fission in the DVC of HFD-fed rats (by expressing a dominant negative form of Drp1, Drp1-K38A, Fig. 1B and Filippi *et al*, 2017) was sufficient to maintain insulin sensitivity and decrease food intake, body weight and fat deposition (Fig. 1G to L). More specifically, a single injection of insulin in the DVC was unable to decrease food intake in HFD-fed rats, while expression of the catalytically inactive form of Drp1 in the DVC of HFD-fed was sufficient to maintain insulin sensitivity and decrease food intake (Fig. 1G). In addition, chronic inhibition of Drp1 in the DVC (through injection of the catalytically inactive Drp1-K38A) led to a decrease in food intake and body weight (Fig. 1H and I), and decreased total white adipose tissue accumulation (sum of retroperitoneal, epididymal and visceral fat) (Fig. 1L). Overall, these data show that HFD-induced changes in mitochondrial dynamics can affect insulin sensitivity in the DVC and feeding behaviour, and that inhibition of Drp1-dependent mitochondrial fission is protective against these deleterious changes.

### Drp1 activity regulates iNOS levels in DVC

How changes in mitochondrial dynamics can affect insulin sensitivity is still unclear. Interestingly, there is a clear increase of iNOS levels in the NTS of the DVC in HFD-fed rats compared with RC-fed rats (Fig. 2A, Fig. S1 and Filippi *et al*, 2017). Therefore, we investigated whether changes in mitochondrial dynamics can affect iNOS levels. Indeed, in the rat neuronal cell line PC12, expression of the active form of Drp1 (Drp1-S637A) led to an increase of iNOS levels (Fig. 2B) when compared to the expression of the dominant negative form of Drp1 (Drp1-K38A) or GFP (Fig. 2B). Changes in iNOS levels in the brain can cause an associated increase in nitric oxide (NO) levels. NO can act as a neurotransmitter or is used to nitrosylate cysteine (S-nitrosylation) or tyrosine (Nytril-tyrosine) residues in a plethora of signalling molecules, thus changing their activity (Mehta *et al*, 2008; Lee & Choy, 2013; Yang *et al*, 2015; Suárez *et al*, 2009). We analysed the S-nitrosylation levels of proteins upon expression of Drp1-S637A, Drp1-K38A and GFP in PC12 cells, and as predicted, cell lysates expressing Drp1-S637A to increase mitochondrial fission exhibited an increase in S-nitrosylated proteins when compared with cell lysates expressing Drp1-K38A or GFP (Fig. 2C). Nitrosylated protein levels were decreased in PC12 cells when iNOS was knocked down using a lentiviral system expressing shRNA for iNOS (shiNOS), confirming that changes in iNOS levels can alter the nitrosylation pattern in PC12 cells (Fig. 2D and Fig. S2). Thus, the changes in nitrosylation levels could be a consequence of the higher iNOS expression levels in PC12 cells expressing Drp1-S367A. Taken together our results show that increasing mitochondrial fission is sufficient to increase iNOS levels and consequently increase S-nitrosylated proteins in PC12 neuronal cells. We then investigated whether changes in mitochondrial dynamics can affect iNOS levels in vivo. Expression of the active form of Drp1 in the DVC of RC-fed rats was sufficient to increase iNOS levels ∼2.5 times while expression of the inactive form of Drp1 decreased iNOS levels ∼ 36% in the DVC of HFD-fed rats (Fig. 2E and F). These data were also confirmed by IF; in RC-fed rats, NTS areas expressing Drp1-S637A presented higher iNOS protein levels compared with areas of the DVC where Drp1-S637A was not expressed (Fig. 2G). On the other hand, in HFD-fed rats, the NTS areas expressing the inactive form of Drp1 presented lower iNOS protein levels when compared with other DVC areas where Drp1-K38A was not expressed (Fig. 2H). Altogether our findings suggest that changes in iNOS levels caused by altered mitochondrial dynamics could be involved in the development of insulin resistance in the DVC.

**Figure 2:**
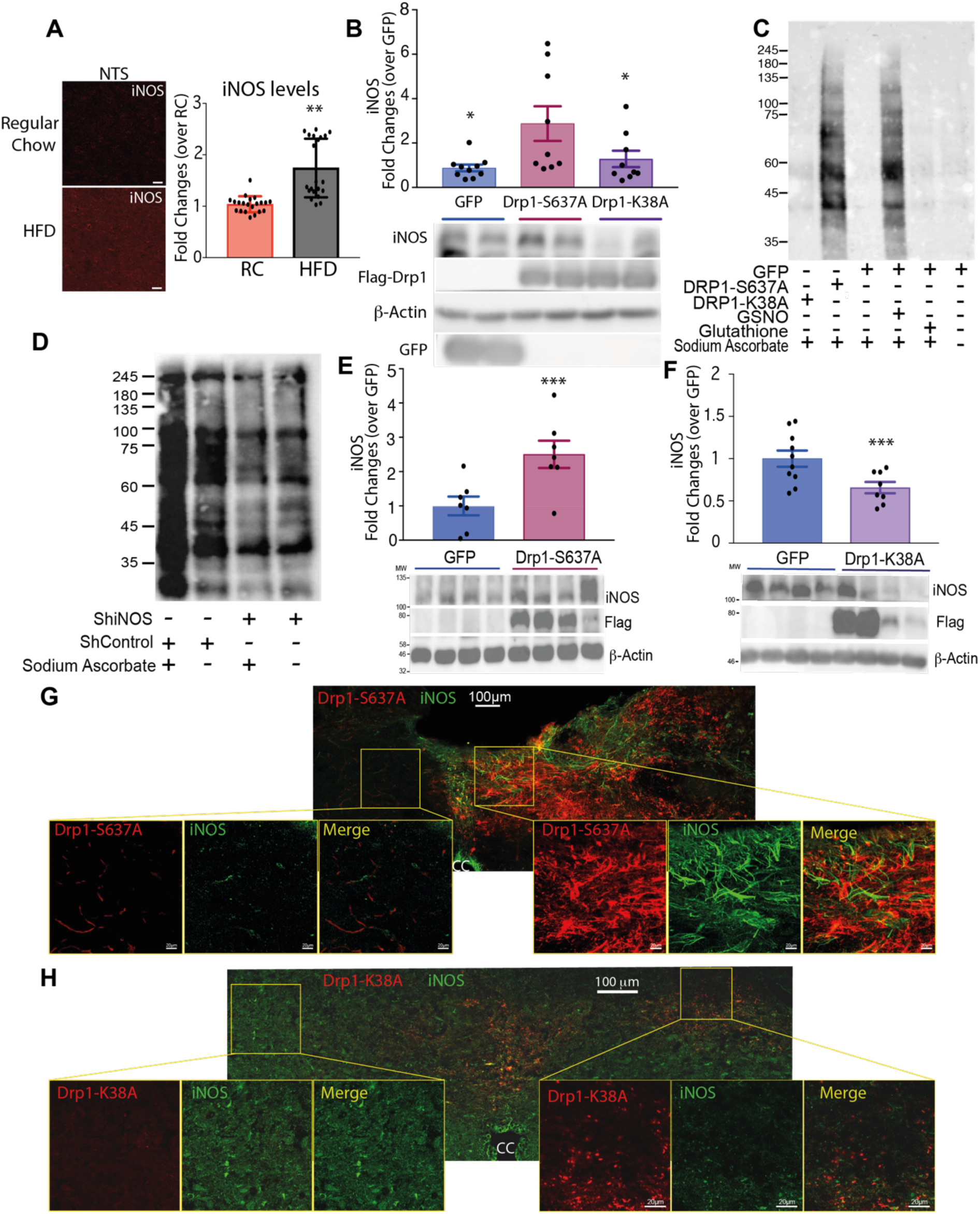
Activation of Drp1 induces an increase in nitrosylated proteins and iNOS, while inhibition of Drp1 in HFD-fed rats decreases iNOS levels. **(A)** Immunofluorescence (IF) representative images (see also Fig. S1) with quantification of iNOS levels in the NTS of rats fed with RC or HFD. Data are average of 21 images taken from, 2 RC or 2 HFD-fed rats. Bar=20μm **(B)** iNOS levels in PC12 cells expressing GFP, Drp1-S637A or Drp1-K38A. **(C)** Nitrosylation levels in PC12 cells expressing either GFP, Drp1-S637A or Drp1-K38A. GFP expressing cells were treated also with 200 mM of S-nitroglutathione or glutathione as a positive and negative control, respectively. Samples were reduced using sodium ascorbate to enable specific labelling of nitrosylated proteins with iodoTMTzero. Sample without sodium ascorbate treatment showing non-specific staining **(D)** Nitrosylation levels of PC12 knocked down for iNOS compared with control PC12 cells (see also Fig. S2). Sample without sodium ascorbate treatment showing non-specific staining **(E)** Western blot analysis of the changes in iNOS levels in RC-fed animals expressing GFP or Drp1-S637A in the DVC (same rats used in Fig 1 C to F). Data are shown as mean ± SEM, with each single point highlighted of n=8 rats for both GFP and Drp1-S637A. **(F)** Western blot analysis of the changes in iNOS levels in HFD-fed animals expressing GFP or Drp1-K38A in the DVC (same rats used in Fig. 1 G to L). Data are shown as mean ± SEM, with each single point highlighted of n=8 for both GFP and Drp1-K38A. **(G-H)** IF representative images of iNOS levels in the NTS of RC-fed rats expressing Drp1-S637A (G) or HFD-fed rats expressing Drp1-K38A (H). *p<0.05, **p<0.01 ***p<0.001

### iNOS knockdown in the DVC prevents HFD-dependent insulin resistance, hyperphagia and body weight gain

Our data indicate that increasing mitochondrial fission in the DVC is sufficient to cause elevated iNOS levels. Previous studies have shown that increased iNOS levels can cause hypothalamic insulin resistance and obesity in mice (Katashima *et al*, 2017). We hypothesised that increased iNOS levels and a consequent increase in S-nitrosylation could be the link between mitochondrial fission and insulin resistance in the DVC. To investigate this possibility, we knocked down iNOS expression in the DVC by injecting a lentivirus expressing shiNOS (Fig. 3A). Control rats were injected with a virus expressing scrambled shRNA (shControl). Using this approach, iNOS levels in the DVC were decreased by 50% (Fig. 3B and C). In order to see whether changes in iNOS protein levels can affect the NO levels in the DVC, we collected DVC wedges from rats infected either with the shiNOS-expressing virus or with the shControl-expressing virus and fed either with RC or HFD. NO is a volatile compound that is hard to measure in tissues, however a reliable estimation can be achieved by measuring NO metabolites like nitrates (Csonka *et al*, 2015). We confirmed that knockdown of iNOS in the DVC led to a significant decrease in nitrate levels thus suggesting that our approach was effective in decreasing both iNOS levels and activity (Fig. 3D).

**Figure 3:**
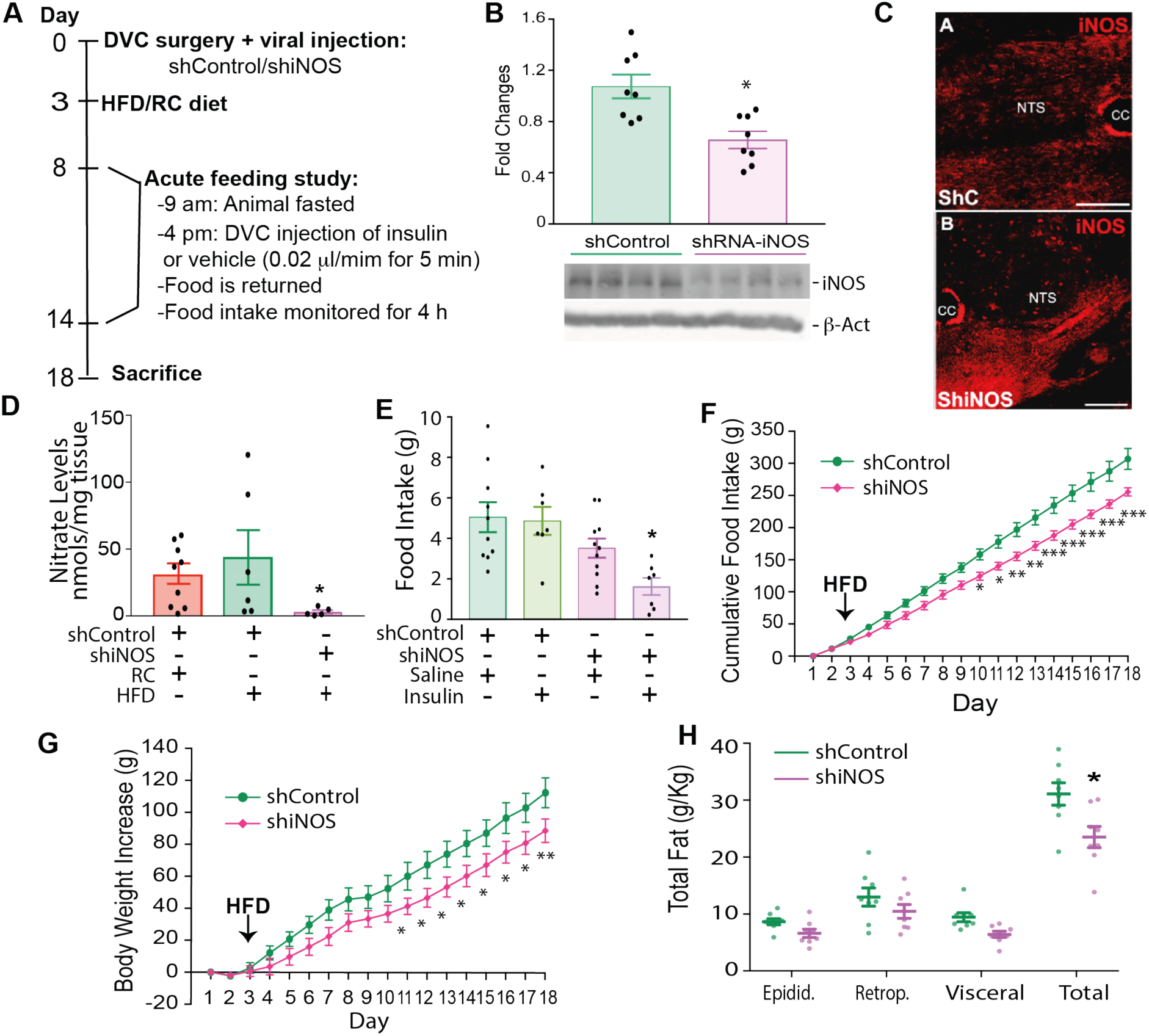
Knockdown of iNOS in the DVC protects from developing HFD-dependent insulin resistance and decreases body weight and food intake. (**A**) Experimental design and feeding study protocol. (**B**) Western blot analysis of the iNOS knockdown in the DVC. iNOS levels of n=8 for shControl and n=8 shiNOS are shown in the bar graph. Representative western blot image is shown at the bottom. (**C**) Representative confocal image of iNOS labelling in animals expressing ShRNA for iNOS or the shControl in the NTS of the DVC. Bar=20μm (**D**) NO levels in the DVC of RC-fed rats compared with HFD-fed rats expressing the control virus and HFD-fed rats expressing shiNOS. Data are shown as mean ± SEM, with each single point highlighted of n=9 RC rats and n=6 HFD-fed rats expressing either shControl or shiNOS. (**E**) Acute feeding study: total food intake at 4 hours comparing animals treated with insulin or a vehicle in the DVC. Data are shown as min ± SEM, with each single point highlighted of n=10 rats for control vehicle, n=7 control insulin, n=11 for shiNOS vehicle, n=7 for shiNOS insulin. (**F**) Chronic cumulative food intake, from day 1 (see schematic in A). (**G**) Chronic data showing body weight increase from day 1. (**H**) White adipose tissue: epididymal, retroperitoneal and visceral fat collected on the day of sacrifice. Data are shown as min ± SEM, with each single point highlighted. Data from F to H are representative of n=10 for shControl and n=8 for shiNOS. *p < 0.05, **p <0.01, *** p<0.001, ****p<0.0001

We next examined whether iNOS knockdown in the DVC could protect HFD-fed rats from developing insulin resistance. Indeed, HFD-fed rats with decreased levels of iNOS in the DVC responded to a single injection of insulin in the DVC by decreasing food intake while HFD-fed rats expressing the control virus remained insulin resistant (Fig. 3E). In addition, chronic decrease of iNOS levels in HFD-fed rats (within 16 days from viral infection-Fig. 3A), reduced food intake (Fig. 3F), body weight gain (Fig. 3G) and total white adipose tissue accumulation (sum of retroperitoneal, epididymal and visceral fat) (Fig. 3H). In summary decreasing iNOS levels in the DVC is sufficient to prevent insulin resistance and reduce food intake, body weight gain and fat deposition in HFD-fed rats.

### Inhibition of Drp1 or knockdown of iNOS in the DVC restores insulin sensitivity in obese rats

Our data show that Drp1 inhibition or iNOS knockdown in the NTS of the DVC is sufficient to prevent the development of insulin resistance in a short term HFD-fed model where HFD is delivered after the viral injection. This suggests that our experimental approach may also be beneficial in an obese model. Therefore, we determined whether insulin sensitivity in an HFD-fed obese rat model could be restored by treating animals with a molecular inhibitor of Drp1 or by knocking down iNOS in the DVC. We used a well-established obese model where rats are given a 4-week (28-days) HFD (Cote *et al*, 2015; Yue *et al*, 2015; Maurer *et al*, 2017). Twenty-eight days of HFD feeding led to a sustained increase in calorie intake (Fig. 4A) and a ∼10% increase in body weight when compared with RC-fed rats (Fig. 4A). Blood glucose levels were also elevated in obese rats when compared with RC-fed rats (Fig. 4C). On day 28, we performed brain surgery and viral injection. Drp1-K38A or GFP injection was carried out the day after surgery (Fig. S3) and delivery of shiNOS or shControl was done on the day of the surgery (Fig. S3). We first investigated whether inhibition of mitochondrial fission or reduction of iNOS levels in the 28-day HFD-fed model could restore insulin sensitivity by performing a feeding study on days 9 and 14 after surgery (Fig. S3). As expected, infusion of insulin in the DVC of RC-fed control animals (expressing either GFP or shControl in the DVC) successfully decreased food intake while our obese model was insulin resistant (Fig. 4D and E). Interestingly, animals expressing the inactive form of Drp1 (Drp1-K38A), had a reduction in food intake in response to DVC-insulin treatment, which was similar to RC control littermates (Fig. 4D). In a similar way, obese animals expressing the shiNOS vector to knockdown iNOS in the DVC presented with decreased food intake in response to DVC-insulin treatment over a four-hour period, while their obese control littermates exhibited insulin resistance (Fig 4E). Overall, these data suggest that either inhibiting Drp1 or decreasing iNOS protein levels in the DVC is sufficient to restore insulin sensitivity in HFD-fed obese rats.

**Figure 4:**
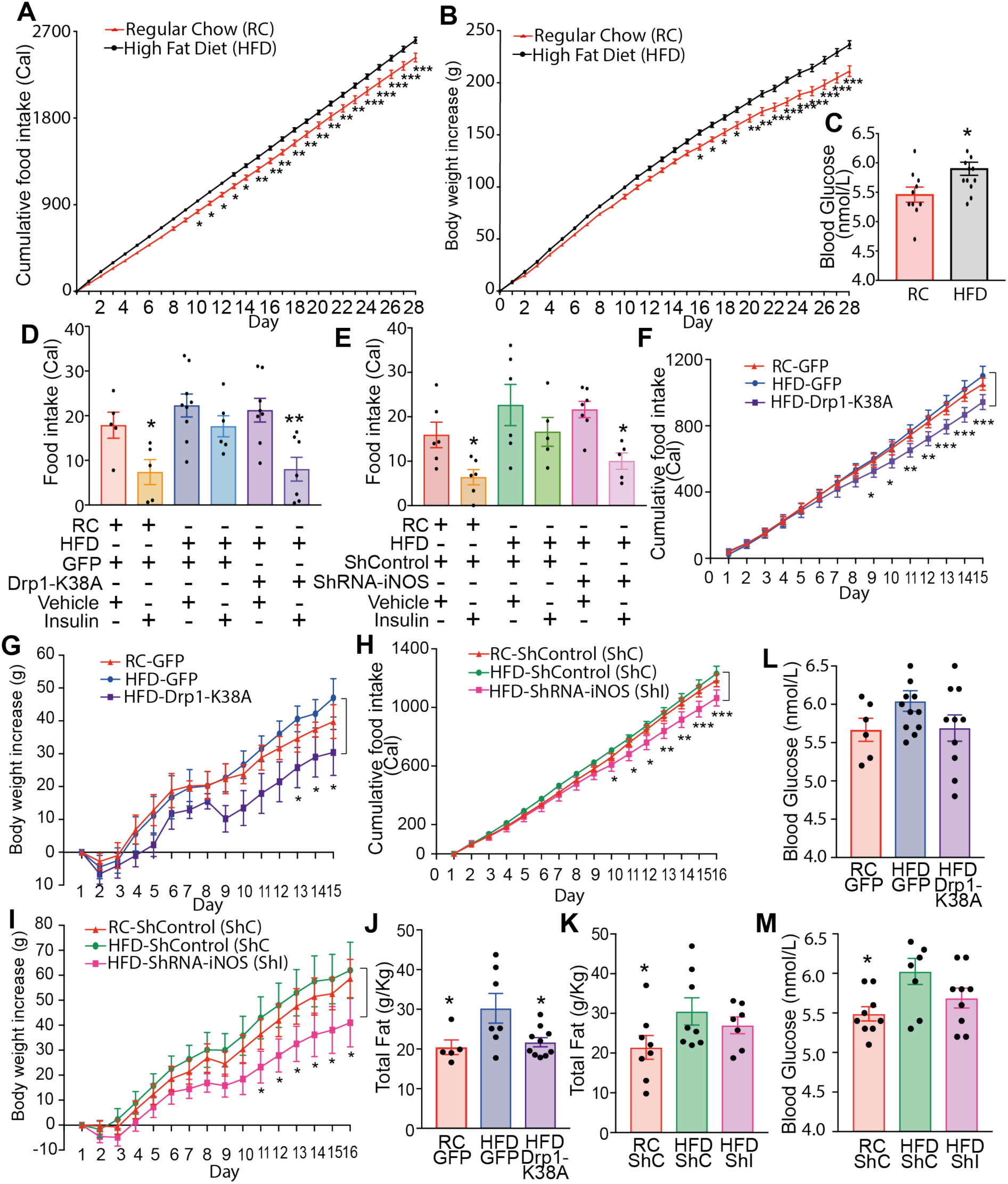
Inhibiting mitochondrial fission and knocking down iNOS in obese rats successfully restored insulin sensitivity and decreased body weight and food intake. Rats were fed for 28 days with HFD or control RC diet. On day 28 rats received DVC surgery. ShControl (ShC) and ShiNOS (ShI) virus were injected on surgery day while the GFP and Drp1-K38A viruses where injected on day 29. Acute feeding study was preformed 8 and 14 days after surgery (see Fig S3). **(A)** Body weight increase over 4-weeks in HFD-fed compared to RC fed animals **(B)** Cumulative food intake over the 4-weeks pre-surgery. Values are multiplied by calories of diet: 3.93Kcal/g RC, 5.51Kcal/g HFD. **(A-B)** Data are expressed as mean ± SEM, n=10 RC, n=24 HFD **(C)** Blood glucose pre-surgery. Bar charts represent mean ± SEM of individual rats shown as single points. **(D and E)** Acute feeding study in GFP and Drp1-K38A expressing animals (D) and ShC and ShI expressing animals (E). Graph shows the total food intake at 4 hours. Bar charts represent mean ± SEM of individual rats shown as single points (n=5 for RC GFP vehicle, n=5 RC GFP insulin, n=9 for HFD GFP vehicle, n=6 for HFD GFP insulin, n=8 for HFD Drp1-K38A vehicle, n=6 for HFD Drp1-K38A insulin; n=6 RC ShC insulin, n=6 for HFD ShCl vehicle, n=5 for HFD ShC insulin, n=7 for HFD ShI vehicle, n=5 for HFD ShI insulin). **(F and H)** Cumulative food intake starting from the day after surgery. **(G and I)** Body weight increase starting from the day after surgery **(G). (J and K)** Total white adipose tissue (sum of epididymal, retroperitoneal and visceral fat) collected on the day of sacrifice. Data are expressed as mean ± SEM, of n=5 GFP RC, n=6 GFP HFD, n=10 Drp1-K38A HFD (F, G and L) and of n=8 ShC RC, n=8 ShC HFD, n=7 ShI HFD (H, I, K). **(L and M)** Average blood glucose over 14 days of the study, data is an average of sampled readings taken before feeding studies and day of sacrifice. Data are expressed as mean ± SEM, n=6 GFP RC, n=15 GFP HFD, n=12 Drp1-K38A HFD and n=9 for ShC RC, ShC HFD, ShI HFD. *p <0.05, **p <0.01, *** p<0.001.

### Chronic Drp1 inhibition or iNOS knockdown in DVC reverses obesity in rats

We then analysed the effect that chronic inhibition of Drp1 or iNOS knockdown in the DVC of obese rats has on food intake, body weight, blood glucose levels and fat deposition. While there was little difference in food intake and body weight between RC-fed and HFD-fed control rats expressing GFP, the rats expressing Drp1-K38A presented with a significant decrease in food intake, associated with a decrease in body weight gain (Fig. 4F and G). We also noted a trend toward a decrease in blood glucose when compared with the HFD-fed rats (Fig. 4L). Similarly, iNOS knockdown caused a decrease in body weight, food intake and blood glucose levels (Fig. 4H, G and M). HFD-fed, obese rats also showed a significant increase in total adipose tissue deposition compared to RC-fed rats (Fig. 5J and K). Drp1 inhibition in the DVC led to a decrease in total fat to the RC-fed level, whereas the iNOS knockdown led to a trend towards fat decrease, although this was not statistically significant (Fig 4J and K).

**Figure 5:**
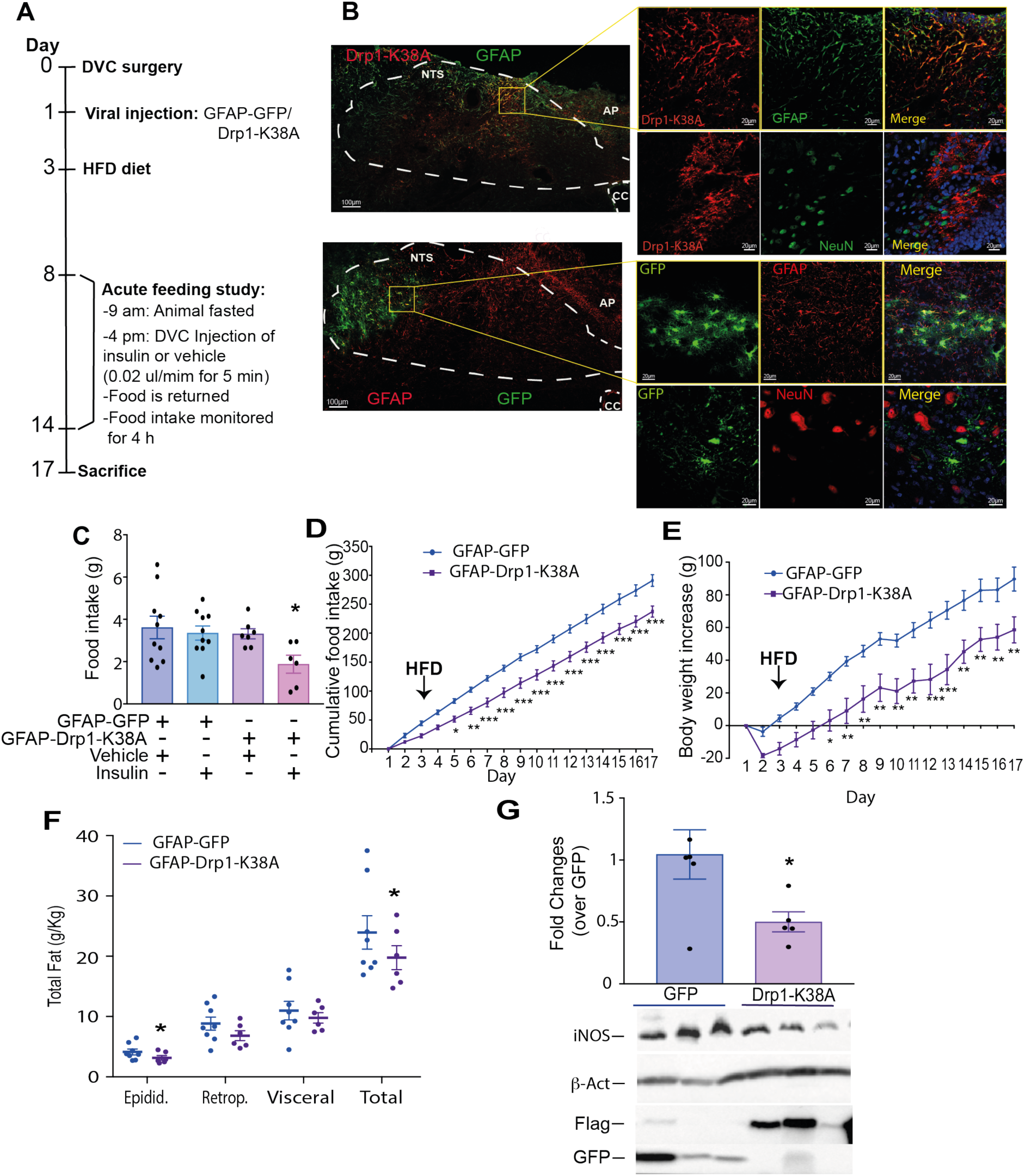
Inhibition of mitochondria fission in astrocytes of the DVC prevent the development of insulin resistance and decreases food intake, body weight gain and fat deposition in HFD-fed rats. (**A**) The experimental design, including the feeding study procedure. (**B**) Representative confocal image of DVC areas expressing FLAG-tagged Drp1-K38A or GFP under the GFAP promoter in the NTS of the DVC. Astrocytes are labelled with anti-GFAP antibody while neurones are labelled with anti-NeuN antibody. Bar=100 μm for the full image and 20 μm for the magnified tiles (**C**) Acute feeding study: animals fed with HFD were fasted for 6 hours and then infused bilaterally into the DVC with a total 0.2 μl of insulin or a vehicle over 5 minutes. Food was then returned and food intake was observed every half hour for 4 hours. Data are shown as min ± SEM, with each single point highlighted. Data are representative of n=10 for GFP vehicle, n=10 for GFP insulin, n=7 for Drp1-K38A vehicle, n=6 for Drp1-K38A insulin (**D**) Cumulative food intake from day of 1 (Fig 5A). (**E**) Body weight increase from day 1. White adipose tissue measurements: epididymal, retroperitoneal and visceral fat collected on the day of sacrifice. Data are shown mean ± SEM, with each single point highlighted. Data in D to F are representative of n=7 rats for both GFP and Drp1-K38A. (**G**) Western blot analysis of changes in iNOS levels in HFD-fed animals expressing GFP or Drp1-K38A in the astrocytes of the DVC. Representative western blot images of iNOS, β-actin, Flag, GFP are also shown. Data are shown as mean ± SEM, with each single point highlighted for n=6 animals expressing GFP and n=5 animals expressing Drp1-K38A. *p < 0.05, **p <0.01, *** p<0.001.

Collectively, our data suggest that inhibition of mitochondrial fission through Drp1 or iNOS knockdown in the DVC can restore insulin sensitivity, decrease body weight gain, food intake and fat deposition in diet-induced obese rats, along with a moderate decrease in glycemia.

### Inhibiting mitochondrial fission in DVC astrocytes of HFD-fed rats ameliorates metabolic health

Our viral injection approach targets multiple cell populations in the NTS by expressing Drp1 under a CMV promoter. The cell population(s) in the DVC which are involved in insulin sensing and insulin resistance are still unknown. We analysed the cellular localization of the GFP expressing adenovirus and observed a high number of GFP-positive astrocytes (∼40 %) followed by neurones (24%) and oligodendrocytes (∼36%) (Fig. S4A-C). A similar pattern of expression was also observed for Drp1-S637A and Drp1-K38A infected cells in the DVC (Fig. S5).

Astrocytes are the most abundant cell types in the brain and they regulate multiple aspects of neuronal function including synaptic plasticity, survival, metabolism and neurotransmission (García-Cáceres *et al*, 2016). Interestingly, astrocytes also synthesise and release NO, which is used for communication with the surrounding neurones (Mehta *et al*, 2008). In addition, chemogenetic activation or inhibition of astrocytes in the DVC was recently reported to affect feeding behaviour (MacDonald *et al*, 2019). These observations led us to hypothesise that altering mitochondrial dynamics in astrocytes may be sufficient to recapitulate the phenotypic changes we observed when targeting all cell types of the DVC.

We tested this hypothesis by using a targeted adenoviral expression system under the control of a GFAP promoter (Yang & Wang, 2015). We generated viruses that express the dominant negative from of Drp1 (GFAP-Drp1-K38A) and a GFP control (GFAP-GFP). On day 0 rodents underwent a stereotactic surgery where a bilateral cannula was inserted into the NTS of DVC. On day one rodents were injected with either a control virus expressing GFAP-GFP or GFAP-Drp1-K38A, and on day three these rodents were given HFD for 14 days (Fig. 5A). To investigate whether we achieved a targeted expression in astrocytes we performed IF, where we confirmed the presence of co-localization between GFAP-GFP or GFAP-Drp1-K38A and an astrocytic marker (GFAP) while there was no co-localization with a neuronal marker (NeuN) (Fig. 5B).

The feeding study was performed on day 8 and 14 (Fig. 5A). Fasted rodents were injected with insulin or a vehicle bilaterally into the DVC, and food intake was measured every 30 minutes for four hours (Fig. 5A). Rodents expressing GFAP-Drp1-K38A had a 37% decrease in food intake in response to insulin compared to GFAP-GFP expressing rodents that were insulin resistant (Fig. 5C). Chronic inhibition of Drp1 in the astrocytes of the DVC produced a clear decrease in food intake and body weight (Fig. 5D and E). Fat deposition was also decreased in rats expressing Drp1-K38A in astrocytes when compared with GFP-expressing rats (Fig. 5F). These data show that inhibition of Drp1 in astrocytes of the DVC is sufficient to prevent HFD-dependent development of insulin resistance, hyperphagia, body weight gain and fat accumulation. At the molecular level there was a reduction of iNOS levels in the DVC of HFD-fed rats expressing GFAP-Drp1-K38A when compared with GFAP-GFP expressing rats.

Interestingly the effect on food intake and body weight was apparent from day 2, when rats where still fed with RC diet, these data suggested that also in a RC-fed rodent inhibition of mitochondrial fission could have an effect on feeding behaviour. To confirm our hypothesis, we performed a feeding study in RC-fed rats expressing either Drp1-K38A or GFP in astrocytes and we could confirm that Drp1-K38A expression in astrocytes decreased both food intake (Fig. S6A) and body weight (Fig. S6B) while no differences in fat deposition where observed (Fig. S6C).

In summary, modulation of mitochondrial dynamics in astrocytes of the DVC is sufficient to regulate iNOS levels, insulin sensitivity, feeding behaviour, body weight and fat deposition in HFD-fed rats.

## Discussion

We have shown that increasing DVC mitochondrial fission triggers insulin resistance, causes hyperphagia, increases body weight gain and fat deposition (Fig 6a). Conversely, inhibiting DVC mitochondrial fission protects rats from developing HFD-dependent insulin resistance, hyperphagia and body weight gain (Fig 6 b). Using a 28-day HFD-fed obese model, we also demonstrated that inhibition of mitochondrial fission restores insulin sensitivity and decreases body weight and fat deposition after a long term HFD-feeding (Fig 6 e). Furthermore, we provide evidence that increased DVC mitochondrial fission leads to higher iNOS levels. Knocking down iNOS in the DVC prevents insulin resistance in HFD-fed rats and reduces hyperphagia and body weight gain (Fig 6 c, f). Finally, we show that inhibiting mitochondrial fission in astrocytes is sufficient to protect HFD-fed rats from developing insulin resistance and results in lower food intake, lower body weight and fat deposition (Fig. 6 d).

**Figure 6:**
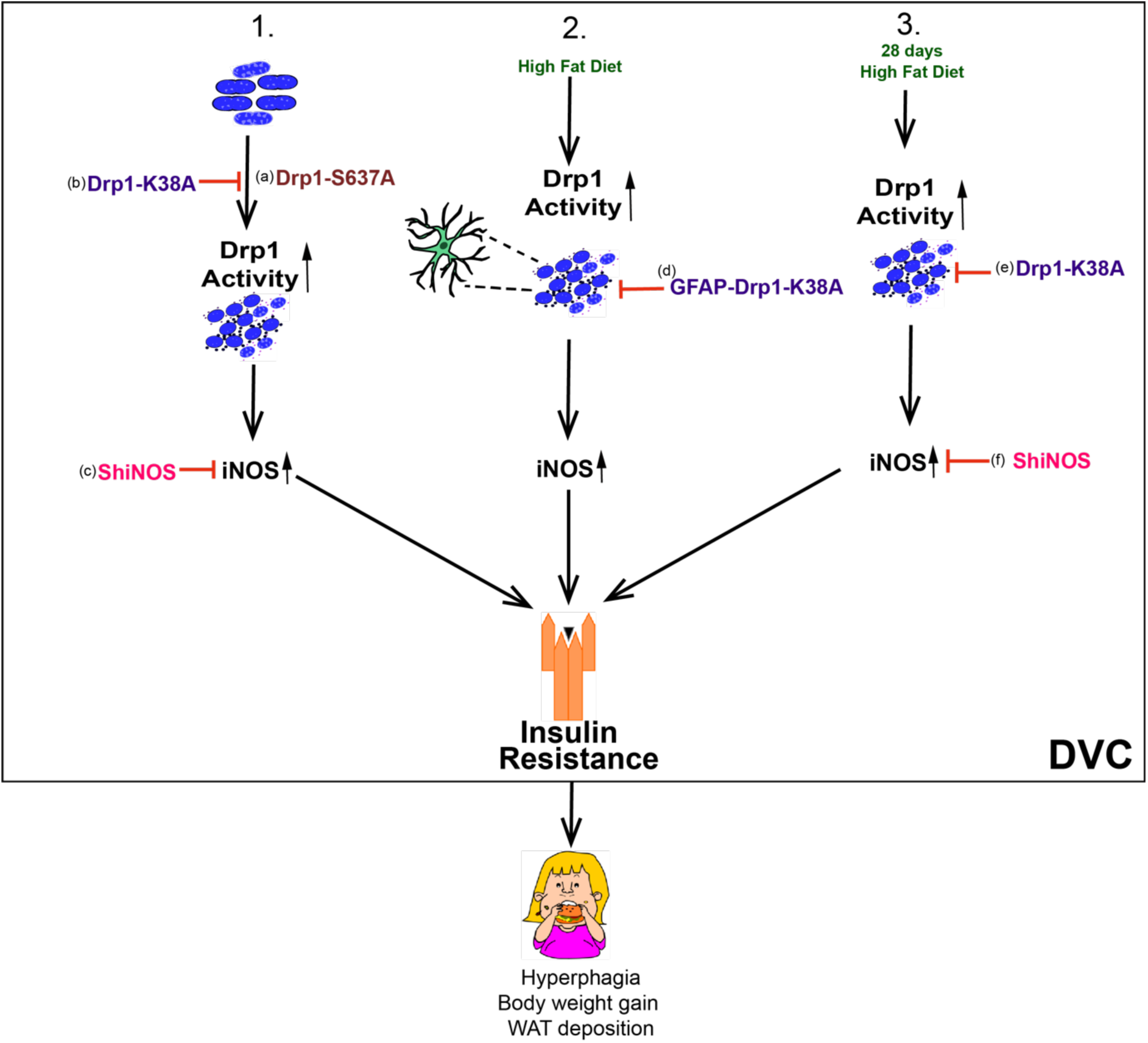
Summary of Drp1 regulation of insulin resistance and body weight. (1) Activation of Drp1 in the DVC causes insulin resistance, hyperphagia and body weight gain (a) while inhibition of Drp1 (b) or decreasing iNOS levels (c) in the DVC is sufficient to protect from HFD-dependent insulin resistance, hyperphagia and body weight gain. (2) Inhibition of mitochondria fission in DVC astrocytes by expressing Drp1-K38A (d), protects HFD-fed rats from developing insulin resistance and lowers food intake, body weight and fat deposition. (3) Inhibition of mitochondria fission (e) or knockdown of iNOS (f) can restore insulin sensitivity and decrease body weight gain and fat deposition in a 28-day HFD-fed obese model.

Our findings are consistent with studies demonstrating that alterations to mitochondrial dynamics in both anorexigenic (POMC) and orexigenic (AgRP) neuronal subtypes within the hypothalamus can affect feeding behaviours and glucose metabolism. Indeed, deletion of Drp1 in POMC neurons improves glucose metabolism (Santoro *et al*, 2017). Additionally, loss of Mfn-1 or Mfn-2 in POMC neurones leads to impaired glucose sensing and insulin release, defective insulin sensing and an increase in reactive oxygen species (ROS) and ER-stress (Ramírez *et al*, 2017; Schneeberger *et al*, 2013). While inhibition of Mfn-1 and 2 in AgRP neurons in the hypothalamus prevented diet-induced obesity in rodents fed with HFD (Dietrich *et al*, 2015),

Our data shown for the first time that activation of mitochondrial fission in the DVC can induce hyperphagia in RC-fed rodents. Furthermore, inhibition of Drp1-dependent mitochondrial fission in the DVC of HFD-fed rodents prevents hyperphagia and body weight gain and restores insulin sensitivity in a diet-induced obesity model. Indeed, mitochondrial dynamics governed by Drp1 also regulates energy metabolism in other tissues, for instance, insulin sensitivity and mitochondria fragmentation in muscle causes insulin resistance and metabolic imbalance in HFD-fed and obese rodent models (Touvier *et al*, 2015; Bratic & Trifunovic, 2010; Sergi *et al*, 2019). Thus, the mechanistic insights revealed here, are likely to be relevant to other organs and tissues, and may represent a unifying model by which mitochondria regulates glucose metabolism, insulin signalling and body weight.

We showed that Drp1 activation in the DVC leads to an increase in the total amount of WAT deposition in RC-fed rodents, while Drp1 inhibition in HFD-fed rodents resulted in lower WAT deposits. The increase in WAT levels caused by increased mitochondrial fission in the DVC could cause an increase in ROS and FFA levels, potentially exacerbating the insulin resistance by also affecting peripheral organ insulin sensitivity. In fact, an increase in free fatty acids (FFA) increases the oxidation of adipose tissue leading to an accumulation of lipids and mitochondrial dysfunction, this can increase oxidative stress ROS and inflammation in tissues, which is associated with insulin resistance and body weight gain (Slawik & Vidal-Puig, 2006; Gao *et al*, 2014).

Consistent with the fact that HFD causes an increase in iNOS levels in the DVC (Filippi *et al*, 2017), we demonstrated that mitochondria fission increases iNOS levels in the DVC and a reduction of iNOS levels can protect from the development of HFD-dependent insulin resistance and can restore insulin sensitivity in an obese model. iNOS is a marker of inflammation (Fujimoto *et al*, 2005; Soskić *et al*, 2011), where HFD-fed rodents exhibit higher levels of iNOS in muscle leading to insulin resistance (Perreault & Marette, 2001). In addition, hypothalamic infusion of NO, to mimic high iNOS activity triggered insulin resistance and increased food intake (Katashima *et al*, 2017). Interestingly, mRNA levels of iNOS were increased in LPS-treated BV2 microglial cells, however, this phenotype was reversed by treatment with a Drp1 inhibitor, MDIVI-1 (Park *et al*, 2013). Altogether these data indicate that iNOS levels are affected with changes in mitochondrial dynamics and this in turn can alter insulin sensitivity.

How changes in iNOS levels can trigger insulin resistance and reduce body weight and food intake is still not fully understood. We observed that shRNA-mediated knockdown of iNOS significantly reduced NO levels in the DVC of HFD-fed rats. In addition, in vitro we could demonstrate that activation of Drp1 can trigger an increase in iNOS levels and nitrosylated proteins (Fig. 2B and C). This is in agreement with studies in other tissues that show S-nitrosylation of insulin signalling-associated molecules in muscle or key ER-stress-associated molecules in the liver can trigger insulin resistance and obesity (Carvalho-Filho *et al*, 2005; Sugita *et al*, 2005; Perreault & Marette, 2001; Yasukawa *et al*, 2005). A marked increase in iNOS levels was also reported in mouse models of diabetes, while iNOS knockout in HFD-fed mice led to improved glucose tolerance and insulin sensitivity in skeletal muscle (Perreault & Marette, 2001). Interestingly, infusion of NO or S-nitrosoglutathione into the hypothalamus resulted in insulin resistance and an increase in food intake (Katashima *et al*, 2017). Drp1 can also be S-nitrosylated under high oxidative stress which can increase the rate of mitochondrial fission and cause changes in energy balance (Nakamura *et al*, 2010).Therefore, a potential molecular mechanism that triggers insulin resistance in the DVC could be associated with increased iNOS levels and increased nitrosylation of key molecules involved in the transduction of insulin signalling. Indeed, chronic reduction of iNOS levels in the DVC could prevent hyperphagia and body weight gain, by mitigating the HFD-dependent nitrosative stress, and S-nitrosylation of key players in the insulin signalling pathway.

Targeting mitochondrial dynamics in the brain could have beneficial effects in obese subjects. Animal models of obesity exhibit altered mitochondrial dynamics, for example, Zucker rodents display a decrease in mitochondrial functionality which correlated with a decrease in ATP production and an increase in Drp1 activity (Raza *et al*, 2015). Increased calorie consumption also caused impairment of mitochondrial function in the brain, and as a result led to increased ROS production and UPR activation (Pratchayasakul *et al*, 2015; Pipatpiboon *et al*, 2012). We show that inhibiting Drp1 in the DVC of 28-day HFD-fed obese rats is sufficient to restore insulin sensitivity and to decrease food intake and body weight. Notably, recovery after surgery was faster for RC-fed rats when compared with HFD-fed rats, for this reason in the first week post-surgery there was minimal difference in body weight between RC and HFD-fed control rats (Fig. 4G and I). We also saw that decreasing iNOS levels in the DVC has beneficial effects on obese rats. Interestingly, a whole body iNOS knockout model presented with improved glucose homeostasis and increased sensitivity to insulin than their obese control counterparts (Perreault & Marette, 2001). While treatment with an iNOS inhibitor, L-NIL, by ICV injection the ARC of the hypothalamus inhibited HFD-induced astrogliosis, suggesting that iNOS activation may induce hypothalamic inflammation (Lee *et al*, 2018). Therefore, our study strengthens the notion that iNOS may be a good target to ameliorate obese phenotypes.

A major question is which cell types in the DVC are responsible for insulin sensing and resistance. Using a specific targeting of cell types, we demonstrated that inhibition of mitochondrial fission in the astrocytes of the DVC is sufficient to reduce food intake and body weight gain both in HFD-fed and RC-fed rodents. Astrocytes provide metabolic and structural support to neurones and play an active role in neurotransmission. Changes in astrocyte number, density and activity have been associated with HFD-feeding, insulin resistance and obesity (Buskila & Amitai, 2010; Balland & Cowley, 2017). Interestingly, astrocytes in the NTS of the DVC responded to acute nutritional overload by increasing their network complexity to integrate peripheral satiety signals to decrease food intake (MacDonald *et al*, 2019). We could therefore hypothesise that there is a potential neuronal-glial cross talk in which astrocytes sense changes in nutritional levels and affect the way in which surrounding neurones control energy homeostasis, potentially *via* NO release or altering the activity of key signalling molecules *via* nitrosylation.

Interestingly, we have also shown that inhibition of mitochondrial fission in astrocytes of the DVC has an important protective effect on the development of HFD-dependent insulin resistance. Indeed, in the hypothalamus, ablation of insulin receptors in astrocytes affected glial morphology and mitochondrial function and reduced the activation of POMC neurones (García-Cáceres *et al*, 2016). Whether changes in mitochondrial fission in astrocytes of the DVC affect the way in which astrocytes sense insulin or the way in which neurones respond to changes in insulin levels is an important question that warrants further investigation.

The DVC is an underappreciated area of the brain that integrates peripheral cues from the metabolic status of an individual and relays to the forebrain to control and maintain energy balance. Here, we shed light on a complex molecular mechanism that triggers insulin resistance in the DVC to cause metabolic imbalance and highlight how the astrocytes, as energy sensors in the brain, play a major role in this process.

Our work uncovers new molecular and cellular targets that can be exploited to develop new approaches focusing on the brain to counteract the deleterious effect of obesity and diabetes..

## Material and Methods

### Animals

Nine-week-old male Sprague-Dawley (SD) rats weighing between 270-300g (Charles River Laboratories) were used in line with the UK animals (Scientific Procedures) Act 1986 and ethical standards set out by the University of Leeds Ethical Review Committee. Animals were housed individually and maintained on a 12-hour light-dark cycle with access to either regular chow (RC) or high fat diet (HFD) and water ad libitum.

Rats were stereotactically implanted with a bilateral catheter targeting the nucleus of the solitary tract (NTS) within the DVC, 0.0mm on occipital crest, 0.4mm lateral to the midline and 7.9mm below skull surface (Filippi *et al*, 2017) on day 0. On day 0 a lentiviral system was used to deliver ShRNA to knockdown of iNOS (shiNOS) or a control scramble ShRNA (shControl) (Santa Cruz Biotechnology, sc-29417-V and sc-108080 respectively) or on day 1 an adenoviral system was used to deliver either a constitutively active form of Drp1 (Drp1-S637A); a catalytically inactive form of Drp1 (Drp1-K38A) or a control of GFP expressed under CMV (Filippi *et al*, 2017) or GFAP promoters. In the first cohort, animals expressing Drp1-S367A or GFP were given RC. In the second and third cohort, animals expressing Drp1-K38A and GFP or shiNOS and shControl, were given RC for 3 days post-surgery and then were put on HFD. The diet composition for RC (3.93kcal/g) was 20.5% protein, 7.2% fat, 3.5% ash and 61.6% carbohydrate (Datesand group, F4031) or HFD (5.51kcal/g) was 20.5% protein, 36% fat, 3.5% ash, 36.2% carbohydrate (Datesand group, F3282). Animals were sacrificed on day 16 and epididymal, retroperitoneal and visceral fat were collected and weighed. The DVC was collected for western blot analysis or immunohistochemistry. Bromophenol blue was used to confirm that the NTS of the DVC region was successfully targeted in surgery.

### Acute feeding study

On day 8 and day 14 after viral injection (day 1) an acute feeding study was performed. On the morning of the study animals were fasted for 7 hours before the experiment. At 4pm, animals were infused bilaterally with 2mU/μl insulin or a vehicle into the DVC of the brain, a total of 0.2μl was infused over 5 minutes (0.04μl/minute). Food was returned after infusion. Food intake was measured every 30 minutes for 4 hours and then again at 12, 24 and 36 hours.

### Production of recombinant adenoviruses

The preparation of recombinant adenoviruses expressing FLAG-Drp1K38A, FLAG-Drp1S637A and GFP from the CMV promoter were as previously described (Filippi *et al*, 2017) using the RAPAd CMV Adenoviral Expression system (Cell Biolabs, Inc. San Diego, CA, USA.), following the manufacturer’s instructions. To produce vectors expressing these genes under the GFAP promoter, the CMV promoter gene was removed from the pacAD5 CMV shuttle vectors replaced with a rat GFAP promoter gene. Cloning was carried out using standard protocols and DNA extracted and purified using Zymo Research DNA kits (Cambridge Bioscience). The recombinant adenoviruses were amplified in HEK293 AD cells and then purified using a sucrose-based method (Jiang *et al*, 2015). Briefly virus containing media was centrifuged at 3000 rpm for 5 minutes and passed through a 0.4 µm filter. The media was then loaded onto 10% Sucrose/TENS (50mM Tris-Hcl pH7.4, 100mMNacl, 0.5mM EDTA) in a 4:1v/v ratio and centrifuged at 10000 rpm for 4 hrs at 4°C. The resulting virus pellet was resuspended in PBS at 4°C overnight and the purified virus titred using a DAB assay (Adenovirus Monoclonal Antibody (E28H) MA17001 Invitrogen, Vector Lab SK-4100 DAB kit). The titres of the adenoviruses injected in the animals where between 9.5 × 10^8^ pfu/ml and 1.5 × 10^9^ pfu/ml.

### Obese Model

Six-week-old male SD rats weighing between (170-190g) were used, from day 0 these animals were subjected to HFD or a RC for 28 days. The 28 days HFD feeding protocol caused insulin resistance, obesity (∼10 to 15% increase in body weight compared with RC-fed rats) and increased basal insulin and blood glucose levels (Cote *et al*, 2015; Yue *et al*, 2015; Maurer *et al*, 2017). On day 28 animals were implanted with a bilateral catheter targeting the NTS within the DVC. The animals were split into two cohorts, the first cohort was injected with Drp1-K38A or GFP, and the second cohort was injected with shiNOS or shControl. Animals were subjected to the acute feeding study on day 37 and day 41, following the protocol as state above.

### Nitrate measurement

Nitrate levels in DVC brain wedges were measured using the Nitrate assay kit form BioVision (K544) according to the manufacturer instructions. In brief, DVC wedges were weighted and homogenised in nitrate assay buffer. Half of the homogenate was then incubated with Griess reagent 1 and 2 while the other half was incubated with only Griess reagent 1 in order to obtain a blank for each sample. Each blank value was then subtracted from its corresponding sample before calculating the nitrate levels.

### S-nitrosylation TMT-assay

Nitrosylated proteins were detected using a tandem mass tag switch method (Murray *et al*, 2012). Neuronal cell line, PC12 were infected for 48hours with Drp1-S637A, Drp1-K38A or GFP expressing adenovirus. Cell lysates were lysed with HENS buffer (ThermoScientific 90106) GFP lysates were also treated with either glutathione or S-nitrosoglutathione as negative or positive controls, respectively, and left for one hour. A BCA protein assay was used to make up lysates to 1μg/μl in HENS buffer. Unmodified cysteines were then blocked using sulfhydryl-reactive compound (MMTS). S-nitrosylated cysteines were then reduced with sodium ascorbate and specifically labelled with iodoTMTzero. Nitrosylated proteins were determined by SDS page.

### Western blotting

Samples were lysed using a lysis buffer with Pierce Protease Inhibitor tablets, using a homogeniser on ice. Samples were centrifuged at 12,000RPM for 15 minutes at 4°C, supernatants were collected and protein concentration was determined using a Bradford Assay. Proteins were separated using SDS-page (8 or 10%), and transferred at 4°C for 3 hours. Membranes were blocked in 5% BSA in TBST and the primary, Sigma Anti-FLAG M2 F1804 (1:3,000); Aviva System Biology GFP OAE00007 (1:20,000); AbCam Anti-iNOS ab15323 (1:50); Cell Signalling Technology Phospho-PERK 3179 (1:1,000); Cell Signalling Technology Total-PERK 3192 (1:1,000); Cell Signalling Technology β-Actin 3700 (1:50,000);); Invitrogen Anti-TMT 90075 (1:1000) antibody was left to incubate overnight at 4°C. Membranes were imaged using ECL (BioRad Clarity) or (BioRad ChemiDoc™ MP Imaging System). Protein levels were analysed with ImageJ (Fiji).

### Immunofluorescence (IF)

Animals were perfused at the end of the experiment using 4% paraformaldehyde. The brainstem area containing the DVC was taken and frozen in cryoprotectant, 25 μm sections were cut using a cryostat. Sections were labelled for FLAG using Sigma Anti-FLAG M2 F1804 (1:500) and co-stained with markers for neurones (NeuN, 1:2000, Millipore ABN90), astrocytes (GFAP, 1:1000, Abcam ab7260), oligodendrocytes (PanQKI, 1:100, Neuromab 75-168), or microglia (Iba1, 1:1000, Wako 019-19741). To stain iNOS abcam anti-iNOS ab15323 (1:250) was used. Sections were mounted using vectasheild plus 4’,6-diamindino-2-phenylindole (DAPI) to visualise nuclei. Images were taken using Zeiss LSM880 upright confocal laser scanning microscope.

### IF quantification

iNOS intensity staining was quantified with Fiji, by randomly selecting 4 different areas for each picture and measuring the average intensity. For co-localization studies, images were processed and exported using ZEN software, figures are presented as a single plane image. Images were counted using 3 random tiles of each slice, 3 slices were used per rodent, an average of all counts were taken and demonstrated by quantification graphs.

### Statistical Analysis

All data are expressed as the mean ± SEM. Data were analysed using GraphPad Prism 7 software. A significant difference was determined by using a multiple T-tests, one-way ANOVA (post-doc test: Sidak) or a two-way ANOVA (post-doc test: Turkey). N refers to the number of animals used. P < 0.05 was considered to be statistically significant. Significance was defined by: (*) P < 0.05; (**) P < 0.01; (***) P < 0.001; (****) P < 0.0001.

## Acknowledgment

This work was supported by grants from the Royal Society (RG160605), Diabetes UK (2338) Wellcome Trust (UNS63234). BP is supported by a Leeds University Anniversary research scholarship. BMF was supported by a Maire Curie IF (SEP-752408) and an MRC-Career Development Fellowship (MR/S007288/1).

## Author contributions

**B.P**. conducted and designed experiments, performed data analyses and wrote the first manuscript draft. **L**.**N**. assisted with in vivo experiments, performed all histochemistry and participated in the data analysis. **J**.**C**.**G**. assisted with the experiments. **J**.**D**. Contributed in experimental design, discussion of results and critiqued the manuscript. **B**.**M**.**F**. conceived and supervised the project, designed the experiments (and conducted some), and wrote the manuscript.

## Conflict of interest

No conflict of interest

## Notes

### Competing Interest Statement

The authors have declared no competing interest.

