## Supplementary Figures for "Inhibition of Mitochondrial Fission and iNOS in the Dorsal Vagal Complex Protects from Overeating and Weight Gain"

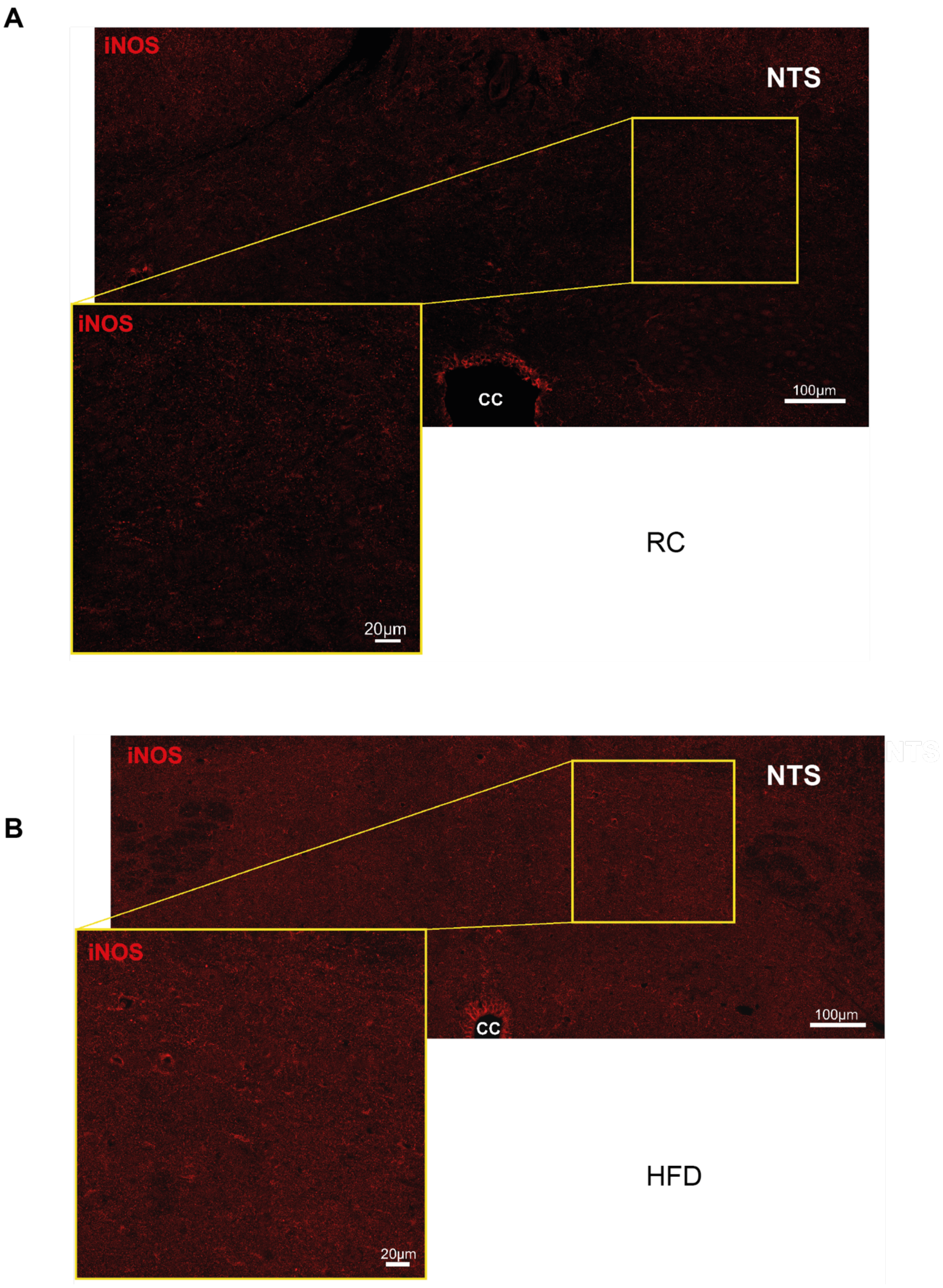


**Figure S1: HFD increases iNOS levels in the DVC.** Representative images of iNOS staining in the DVC of RC (**A**) and HFD-fed (**B**) rats. Image show a large tile that includes central canal and NTS of the DVC. In addition, a magnified area is shown.

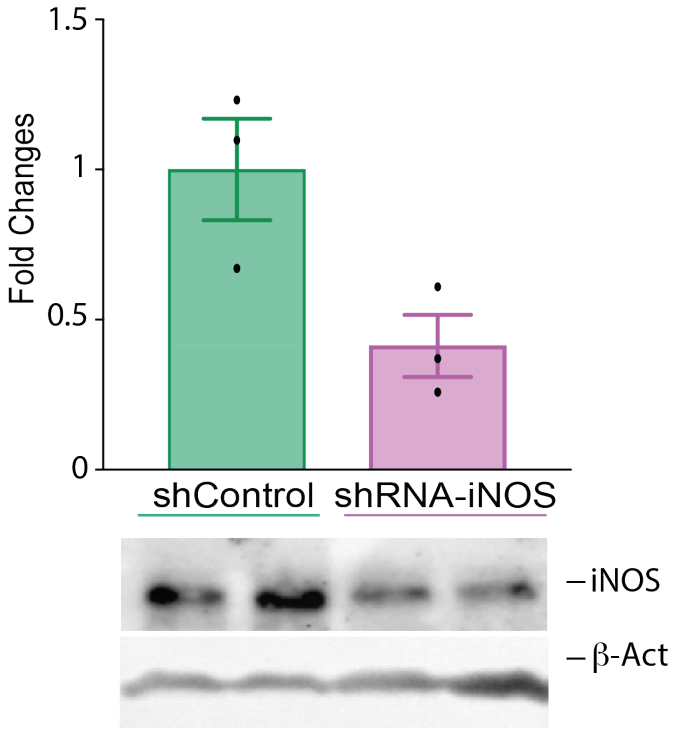


**Figure S2: Control blots for fig. 2 D.** PC12 cells infected with a lentivirus expressing shRNA for iNOS or Control scramble virus where lysed and iNOS knockdown levels quantified by western blotting. In the figure the quantification with a representative western blot is shown. All data are expressed as mean ± SEM *n*=3 for shControl and shiNOS-expressing cells. [*p < 0.05, **p <0.01, *** p<0.001]

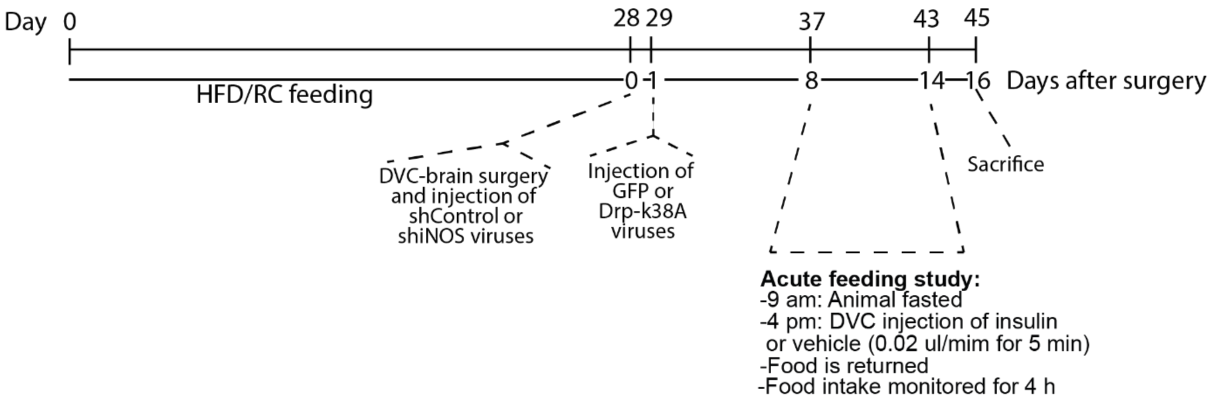


**Figure S3: Experimental design for the 28d HFD-fed obese model**. Rats were fed for 28 days with HFD or control RC diet. On day 28 rats received DVC surgery**.** shControl and shiNOS virus were injected on surgery day while the GFP and Drp1-K38A viruses where injected on day 29. Acute feeding study was performed on day 8 and 14 after surgery.

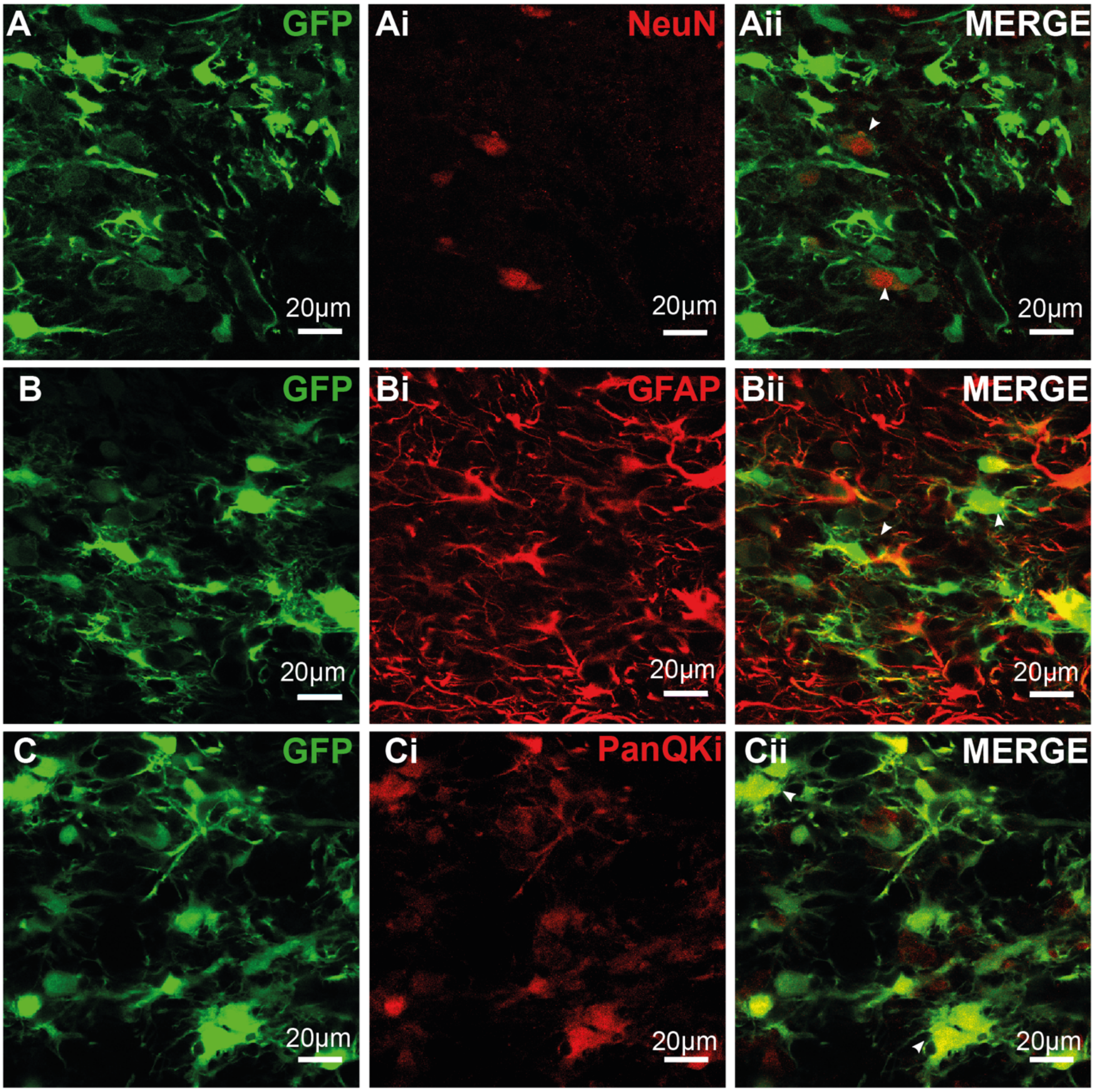


D

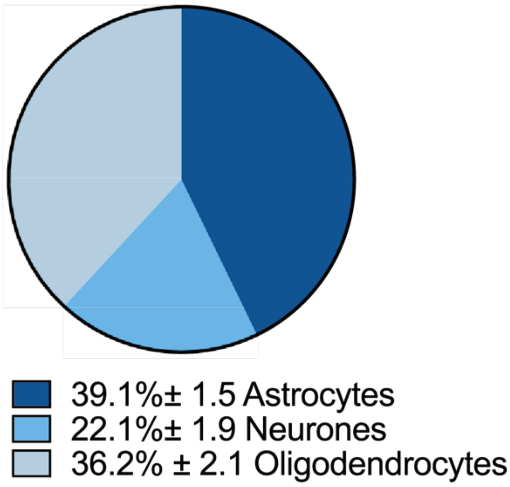


**Figure S4: GFP expression in neural cell types. (A-Aii)** Representative confocal images illustrating labelling of GFP expression in (A), NeuN (Ai) and dual labelling (Aii) in the DVC. **(B-Bii)** Representative confocal images illustrating labelling of GFP expression in (B), GFAP (Bi) and dual labelling (Bii) in the DVC. **(C-Cii)** Representative confocal images illustrating labelling of GFP expression in (C), PanQKi (Ci) and dual labelling (Cii) in the DVC. **(D)** Quantification of the co-localised cells.
Closed arrows denote co-localised cells.
Images and quantification represent the average of n= 3 for NeuN and PanQKi, n= 4 for GFAP animals. 3 images where quantified per animal).

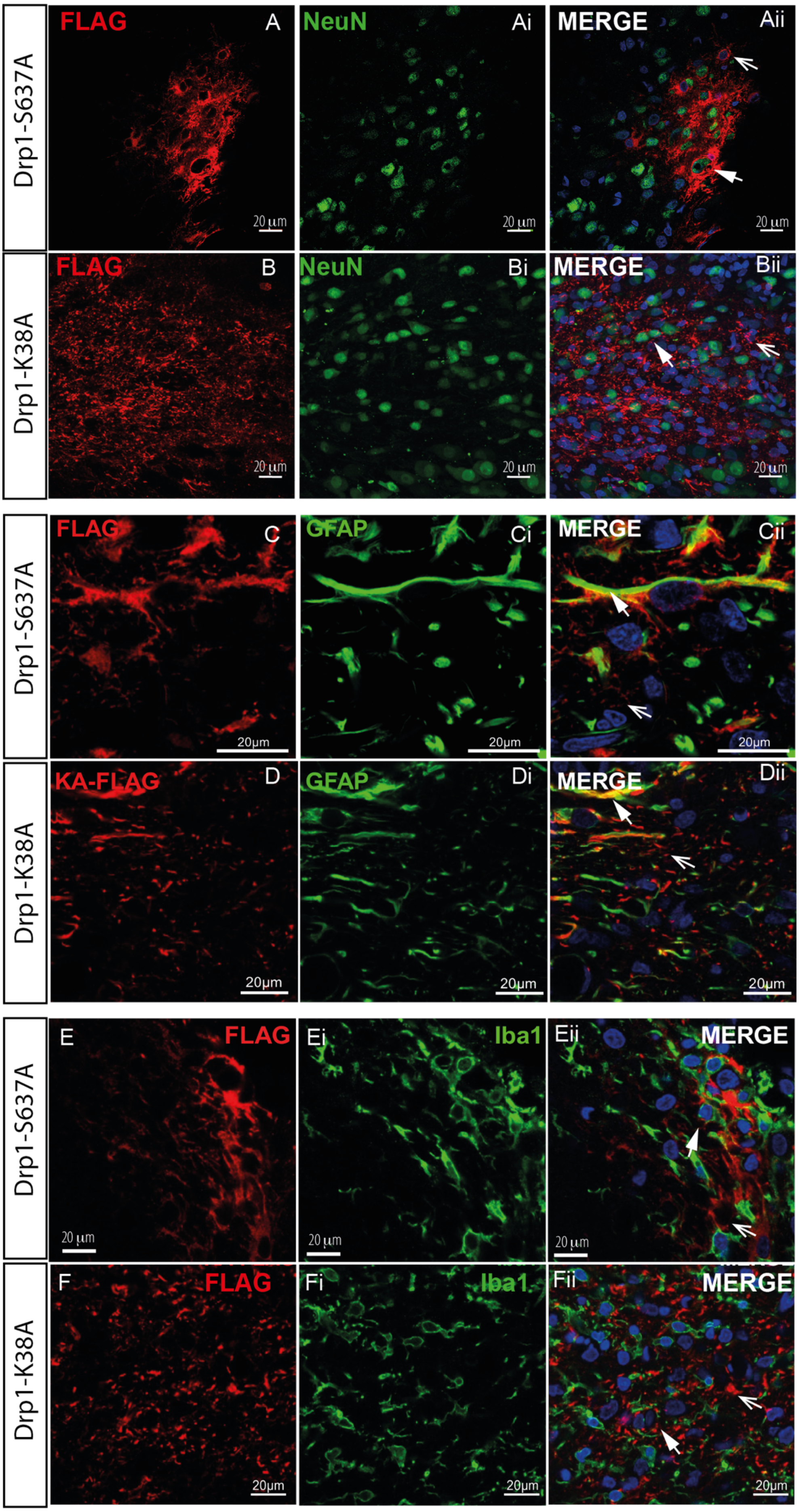


**Figure S5: Expression of the constitutively active form of Drp1, Drp1-S637A and the dominant negative form of Drp1, Dr1-K38A in neuronal cells. (A-Aii)** Representative confocal images illustrating the expression Drp1-S637A (A), NeuN (Ai) and dual labelling (Aii) in the DVC. Blue in Aii represents Nuclei stained with Dapi. **(B-Bii)** Representative maximal intensity confocal images illustrating the expression of Drp1-K38A (B), NeuN (Bi) and dual labelling plus Dapi in blue (Bii) in the DVC. **(C-Cii)** Representative confocal images illustrating the expression of Drp1-S637A (C), GFAP (Ci) and dual labelling plus Dapi in blue (Cii) in the DVC. **(D-Dii)** Representative confocal images illustrating the expression of Drp1-K38A (D), GFAP (Di) and dual labelling plus Dapi in blue (Dii) in the DVC. **(E-Eii)** Representative confocal images illustrating the expression of Drp1-S637A (E), Iba1 (Ei) and dual labelling plus Dapi in blue (Eii) in the DVC. **(F-Fii)** Representative confocal images illustrating the expression Drp1-K38A (F), Iba1 (Fi) and dual labelling plus Dapi in blue (Fii) in the DVC. Open arrows denote non-colocalised cells. Closed arrows denote colocalised cells.

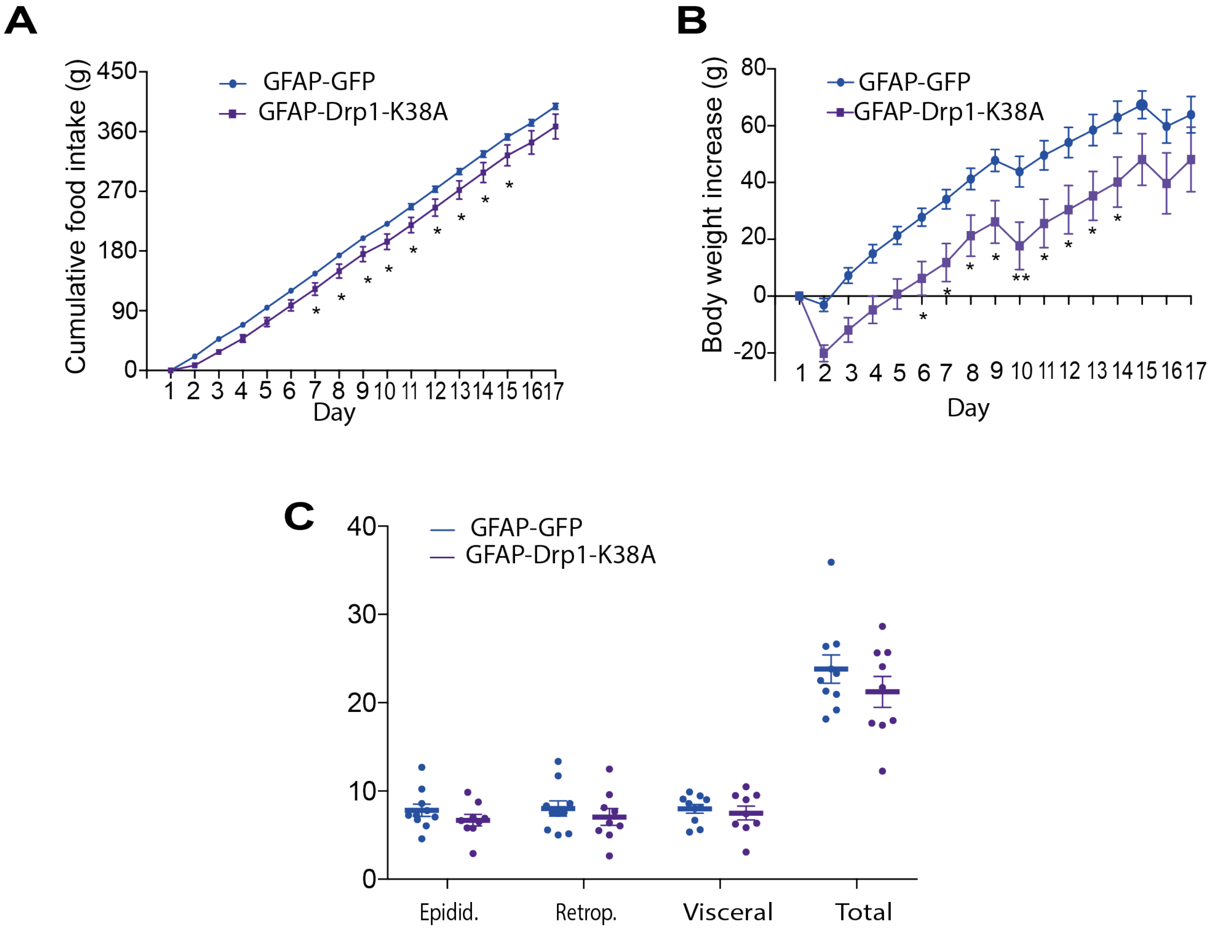


**Figure S6: Inhibition of mitochondria fission in astrocytes of the DVC decreases food intake, and body weight in regular chow-fed rats**. A double cannula was inserted into the NTS of the DVC in RC-fed rats on day 0. On day 1 rats were injected with an adenovirus expressing either GFAP-GFP or GFAP-Drp1K38 in the NTS of the DVC. Food intake and body weight, where measured daily for 17 days. (**A**) Cumulative food intake from day 1. (**B**) Body weight increase from day 1. (**C**) White adipose tissue measurements- epididymal, retroperitoneal and visceral fat collected on the day of sacrifice. Data are shown min ± SEM, with each single point highlighted. Data are representative of n=8 rats for both GFP and Drp1-K38A. *p < 0.05, **p <0.01
